# Lysine Decarboxylation aids in UPEC intracellular survival in the early stages of urinary tract infection

**DOI:** 10.1101/2025.04.25.650641

**Authors:** Michelle A. Wiebe, Seth Reasoner, Tomas Bermudez, John R. Brannon, Grace Morales, Faith Guice, Erin Q. Jennings, Jeffery C. Rathmell, Maria Hadjifrangiskou

## Abstract

Acid stress is a substantial challenge to bacterial life. Acidic conditions can damage the bacterial cell envelope and disturb vital physiological processes, such as enzymatic activity, protein folding, membrane- and DNA maintenance. *Escherichia coli* occupies numerous environmental and host niches with varying pH. Consequently, *E. coli* strains are equipped with multiple acid resistance (AR) mechanisms to withstand acidic conditions. Uropathogenic *E. coli* (UPEC), which accounts for >75% of urinary tract infections (UTIs) persist for years in the host, colonizing the gut and the vagina asymptomatically for long periods of time, while causing acute or chronic infection in the bladder. While these host niches have variable pH, no studies elucidated which AR mechanisms are used by UPEC during infection. Here, we generated a comprehensive list of AR deletion mutants and evaluated them in the acute stage of infection. We show that at the acute infection stage, mutants that lack *cadA* (AR4) are significantly attenuated. We go on to show that AR4 is specifically induced early during intracellular infection within urothelial cells. Deletion of *cadA* leads to fewer intracellular bacteria both *in vitro* and in the murine infection model. Treatment with bafilomycin, which blocks vacuole acidification rescues the *cadA* deletion phenotype intracellularly, suggesting that AR4 is important for UPEC survival inside the lysosome. Combined, this work begins to elucidate the specific contribution of each AR to UPEC pathogenesis.

## Introduction

Extra-intestinal pathotypes of *Escherichia coli*, like uropathogenic *E. coli* (UPEC), cause a global burden of millions of infections annually, most of which are in the urinary tract (1). Urinary tract infections (UTIs) disproportionately affect women, with approximately 80% of all UTIs being reported in women (2). The majority of UPEC strains belong to the B2 and D phylogenetic clades, members of which can colonize the human host for years (3-5). It is hypothesized that UPEC strains can be acquired through ingestion of contaminated food (6), which implies that UPEC can survive the acidic conditions of the stomach and the variable pH encountered in the gastrointestinal and genitourinary tracts (7, 8). In the gastrointestinal tract, stomach acid, bile acids, and organic acid metabolites from the gut microbiota all present pH challenges for UPEC (9-11). In the human vaginal space, where UPEC can form transient asymptomatic reservoirs (12), different *Lactobacillus spp*. (13) excrete large concentrations of lactic acid that lowers the vaginal pH (13-15). In the bladder lumen (**Figure 1A**), the primary immune responders during UPEC infection are neutrophils and macrophages (16), which use mechanisms of bacterial killing that involve low pH. Inside urothelial cells, where UPEC expands during acute infection, the phagolysosome has been shown to kill or expel invading UPEC (16). However, UPEC has been shown to neutralize phagolysosomes in bladder epithelial cells (16, 17). This neutralization prevents bacterial cell death, although the mechanism through which UPEC can accomplish it remains unknown. The pathogenic lifecycle of UPEC in the bladder comprises extracellular and intracellular stages (18). The intracellular expansion during the acute infection stage has been shown in multiple studies to be critical for productive infection, as mutants that are unable to replicate intracellularly have a significant defect in persisting in the host (19, 20). Delineating the specific pathogenic pathways that facilitate UPEC expansion in the urothelial cell are paramount to blocking this critical step in the pathogenic cascade. In previous studies, we reported that UPEC require *de novo* purine biosynthesis during the intracellular stage of acute infection, specifically when inside the vacuole (21). We went on to demonstrate that once inside the host cell cytoplasm, UPEC rely on the Cytochrome bd oxidase for aerobic respiration (19).

**Figure 1.**
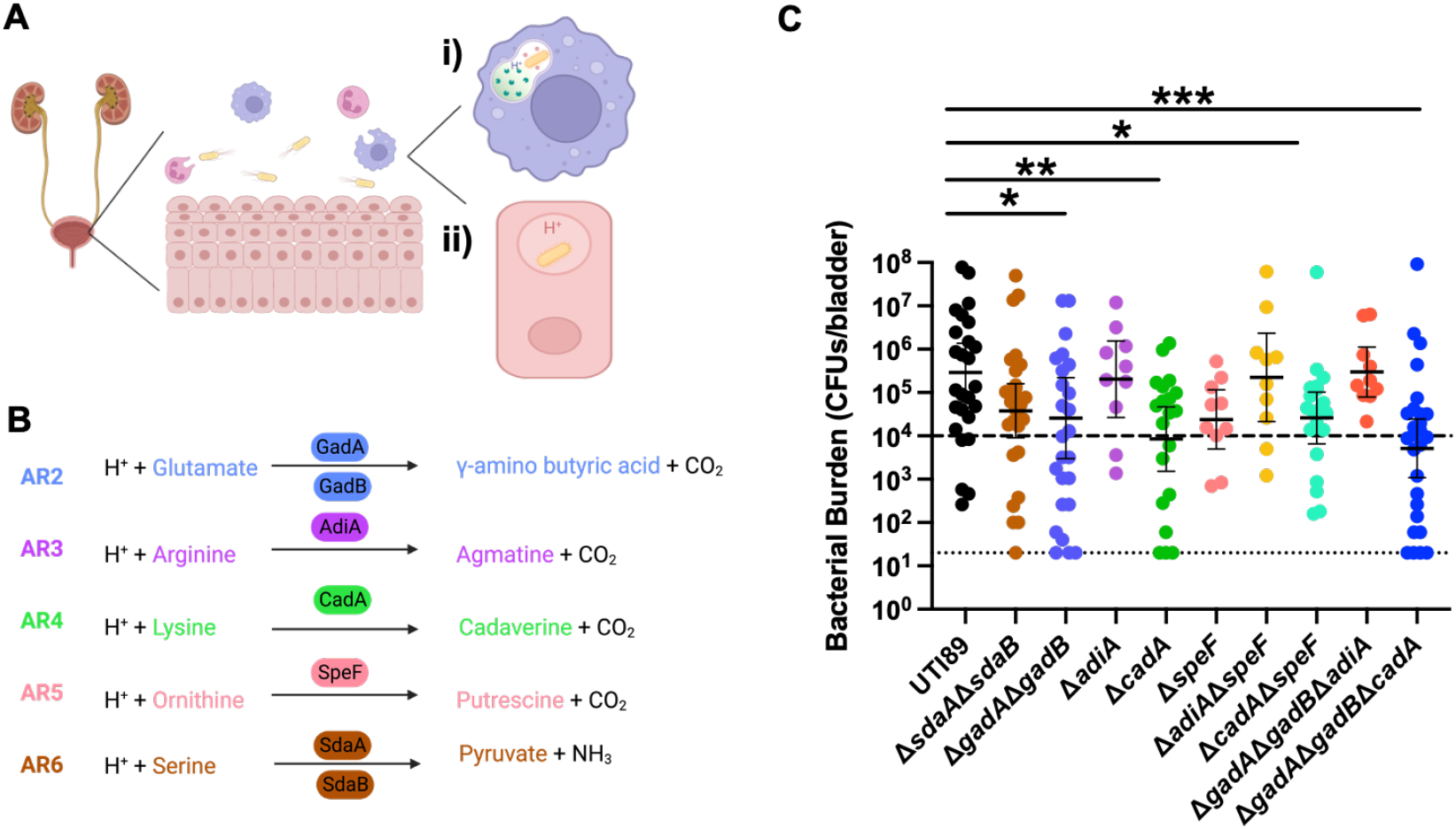
Fitness of Acid Resistance/Response (AR) mechanisms in the bladder. **(A)** Schematic depicts the sources of low-pH stress that UPEC encounters during bladder colonization. Ai) macrophages and neutrophils phagocytose UPEC. Fusion of the phagosome with the lysosome induces a pH decrease in the newly formed compartment. The pH decrease is meant to be bactericidal. Aii) During its pathogenic cascade, UPEC invades bladder epithelial cells. In order to expand into intracellular bladder communities, UPEC must be able to survive and escape from endocytic vesicles that become acidified in response to infection. **(B)** Chemical reactions associated with the different *E. coli* AR systems. AR2-AR5 use decarboxylation reactions that consume a proton to produce CO_2_. The resulting product is then exported by amino acid antiporters. AR6 involves the deamination of serine to produce NH_3_. Schematics in A-B were generated using Biorender. **(C)** Graph summarizes mouse colonization data from a 24h bladder infection. Each dot represents a mouse. C3H/HeN 7-week-old female mice purchased from Envigo were intravesically inoculated with 10^7^ CFUs of bacteria. At 24h post inoculation, mice were euthanized and bacterial titers in the bladder were enumerated. The graph shows the results of two independent experiments. Statistical analysis performed using two-tailed Mann-Whitney. *, p<0.05; **, p<0.001. ***, p<0.0005. Animal work was performed under the approved IACUC animal protocol M1800101-1

In this study, we expand the knowledge into the requirements for intracellular expansion as it pertains to tolerance to low pH while inside the urothelial vacuole. Using a comprehensive panel of strains lacking one or more acid response/resistance (AR) systems (**Figure 1B**), we present evidence that lysine decarboxylation by the CadA/B AR system (AR4) contributes to increased intracellular survival during the early stages of acute UTI.

## Results

### Specific AR systems contribute to bladder colonization in a murine model of bladder infection

The most well characterized AR mechanisms, AR2-5 (**Figure 1B**), increase the cytoplasmic pH by consuming a proton during the decarboxylation of specific amino acids (9, 22-25), while AR6 accomplishes this by de-amination of L-serine and production of ammonium and pyruvate (26). To begin to elucidate the contribution of each AR system in different stages in UPEC pathogenesis, we generated a comprehensive panel of existing (30) and newly created (**Table S1**) AR deletion mutants. Before evaluating the effects of each deletion mutant to adherence and invasion of bladder cells, we determined the effects of each deletion on bacterial growth in laboratory media (lysogeny broth (LB) **Figure S1A**), human urine (**Figure S1B**), or the tissue culture medium used in our studies (RPMI, **Figure S1C**). Growth, measured by OD_600_, in LB is not affected for any of the AR mutants (**Figure S1A**), indicating that deletion of only one or two AR systems is not sufficient to sensitize *E. coli* to changes in pH during *in vitro* growth in rich media. Similarly, during growth in human urine (**Figure S1B**) or RPMI (**Figure S1C**), none of the strains display significant difference in growth compared to the wild-type parent strain, although most strains exhibited a slower growth rate in urine (**Figure S2**). Together, these data indicate that loss of one or two AR systems does not impair growth in the *in vitro* conditions tested here.

Following the *in vitro* analyses, we assessed the ability of each AR deletion strain to colonize the murine bladder. For this work, 7-week-old female C3H/HeN mice were transurethrally inoculated with 10^7^ CFU of wild-type (WT) UTI89, or each of the isogenic mutants. After 24 hours, mice were sacrificed and the bladders, kidneys, and vaginal membranes were harvested, homogenized, and plated for CFUs. Compared to the WT-infected mice, the mice infected with the AR mutants lacking *cadA* (AR4) had significantly lower (1-log) bladder bacterial titers (**Figure 1C**). Similarly, the mutants had decreased titers in the kidney (**Figure S3A**). By contrast, no differences were observed in vaginal titers among strains (**Figure S3B**), suggesting that the AR deletion mutants tested can transit to and colonize the vaginal space, despite having a defect in the bladder.

### AR4 is upregulated during intracellular infection

The acute infection cycle of UPEC encompasses a transient intracellular stage in urothelial cells, where the pathogen replicates in the host cell cytoplasm by consuming oxygen (27). To expand intracellularly, cells must first escape the phagolysosome, where they would encounter a change in pH (**Figure 1A**). We therefore asked whether the defect observed for AR4 mutants *in vivo* stems from an inability to withstand acidification inside the host cell.

To elucidate the contribution of AR4 inside the urothelial cells, we turned to the human urothelial tissue culture model, using the ATCC 5637 (HTB-9) bladder cell line. In this assay, approximately 1% of cells typically become internalized to seed intracellular infection (42). Of these internalized UPEC cells, the majority becomes expelled via a non-lytic mechanism (34, 43), leaving a small portion of cells that drive intracellular expansion. We first asked whether AR4 is specifically upregulated inside urothelial cells. For this experiment, RNA was isolated from intracellular bacteria following invasion of the HTB-9 cell line by wild-type UTI89 (28). Urothelial cells were seeded in 24 well plates and infected with UTI89 at an MOI of 10 and incubated for 2 hours at 37°C with 5% CO_2_. After 2 hours of incubation with the bacteria, wells were treated with gentamicin for 2 hours to kill extracellular bacteria and washed to remove the antibiotic. RNA was extracted from the treated bladder cells, DNAse-treated and reverse-transcribed. Following reverse transcription, transcript abundance of key components of each AR system was determined using probe-based qPCR. Specifically, *gadA, gadC* (AR2), *adiA (*AR3), *cadA* (AR4), *speF* (AR5), *sdaA* and *sdaC* (AR6) transcript abundance was quantified, using the housekeeping gene *gyrB* as a normalizer, as we previously described (27, 29). Transcript abundance intracellularly was compared to the transcript abundance of the same gene in bacterial cDNA prepared from the bacterial input. These analyses revealed that UPEC *cadA* transcript abundance (AR4) is increased in the host intracellular environment, compared to the inoculum (**Figure 2**, Log_2_ fold change = 3.774). Conversely, there was a decrease in *gadC* transcript abundance (AR2) intracellularly (**Figure 2**, Log_2_ fold change = -4.37). The transcripts of the other AR system components remained unchanged, although the *speF* transcript (AR5) appeared to have a nominal, yet not-statistically significant increase (**Figure 2**). These data indicate that UPEC specifically upregulates AR4 in the intracellular environment. We went on to determine the contribution of AR4 to pathogen fitness inside the host cell.

**Figure 2.**
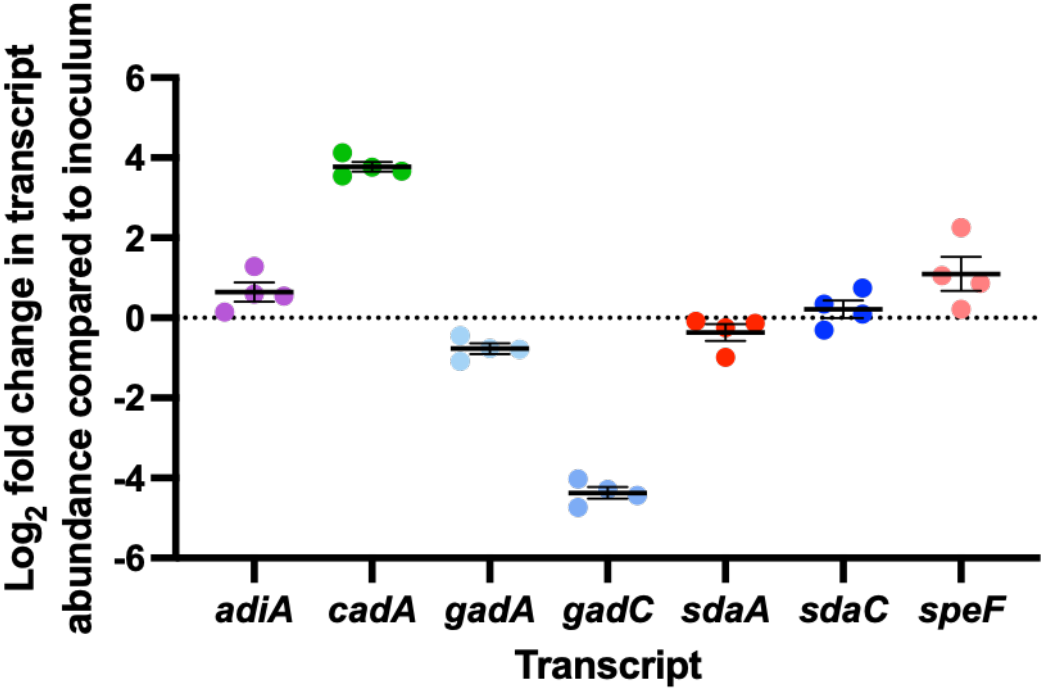
Transcript analysis of AR genes during intracellular bladder cell infection. Graph shows RT-qPCR results of representative genes from each AR system, using RNA isolated from intracellular bacteria. For this work the bladder epithelial cell line 5637 (ATCC HTB-9) was seeded with wild-type UTI89 at an MOI=10. Following a 2h incubation to allow for pathogen internalization, extracellular bacteria were eliminated by incubating the urothelial cell line with gentamicin for another 2 hours. Gentamicin was then removed and cells were washed 3X with 1X PBS, followed by urothelial cell lysis and subsequent treatment for bacterial lysis and RNA extraction. Extracted RNA was DNAse-treated, reverse-transcribed, quantified and subjected to qPCR using TaqMan probes designed for *adiA* (purple), *cadA* (green), *gadA* (pale blue), *gadC* (light blue), *sdaA* (red), *sdaC* (royal blue), and *speF* (coral). Transcript abundance was normalized to the gyrB house keeping gene transcripts. Relative fold changes were determined by the ΔΔ*C*_*T*_ method, relative to the bacterial inoculum corresponding transcripts. Graph shows analysis of 4 biological replicates. Error bars indicate mean and standard error of the mean at 95% CI.

### AR4 mutants are attenuated inside bladder cells

AR4 neutralizes acidic pH by decarboxylating lysine to cadaverine, via the action of the CadA enzyme (**Figure 1B** and (22, 36)). CadB serves as a lysine/cadaverine antiporter. CadA/B genes are regulated by CadC, a membrane integrated transcription regulator **(Figure 3A)**. CadC responds to low pH and changes in lysine abundance to promote transcription of *cadA* and *cadB* (37). LysP, a constitutively expressed lysine importer, has been shown to inhibit *cadBA* expression through interactions with CadC. When extracellular lysine is absent or concentrations are low, LysP inhibits CadC activation of the *cadBA* operon (38).

**Figure 3.**
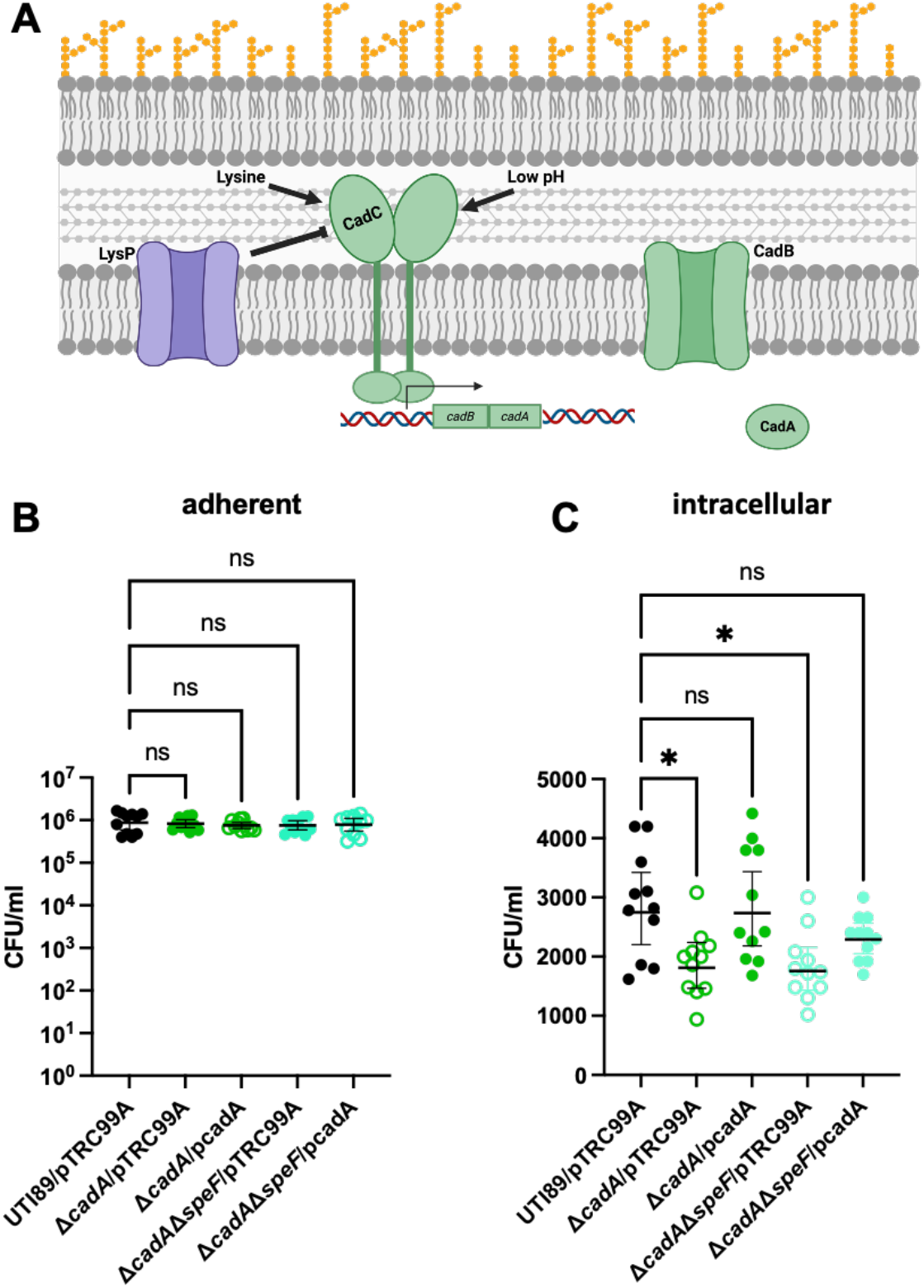
Mutants lacking CadA exhibit a defect in the intracellular compartment. **A)** Schematic depicts the major components of AR4 (Generated using Biorender). Membrane embedded transcriptional regulator, CadC, activates transcription of *cadBA* under high lysine concentration and low pH conditions. LysP, a constitutively expressed lysine importer, inhibits CadC under low lysine concentrations. **B-C)** Graphs show total **(B)** and intracellular **(C)** bacterial titers of wild-type UTI89 (Black), Δ*cadA* (Green), or Δ*cadA*Δ*speF* (cyan) with (filled symbols) and without (open symbols) complementation. Data shown are from two biological repeats. (*p=0.0165, Kruskal Wallis with Dunn’s multiple comparisons test).

To determine if AR4 specifically has defects in the intracellular environment, we inoculated the HTB-9 urothelial cells with wild-type (WT) UTI89, or two mutants lacking cadA: Δ*cadA* (green) and Δ*cadA*Δ*speF* (cyan), given that the pPCR analysis had also demonstrated a nominal increase in the transcript of *speF* (**Figure 2**). Following infection for 2 hours and a 2 hour treatment with gentamicin to kill the extracellular cohort (42), we enumerated total (**Figure S4A**), adherent (**Figure 3B**) and intracellular (**Figure 3C**) bacterial titers. Our analyses revealed that, although total bacterial titers did not differ among strains (**Figure S4A**), in agreement with no apparent growth defects in RPMI media (**Figure S2**), both the Δ*cadA* and the Δ*cadA*Δ*speF* intracellular titers were consistently lower, compared to the WT parent (**Figure 3C**). Complementation of the Δ*cadA* and Δ*cadA*Δ*speF* mutants with a plasmid expressing *cadA* (pCadA) in the backbone of vector pTrc99A, restored intracellular titers of all strains to those of the wild-type strain. Quantification of free lysine inside urothelial cells revealed a consistent, although not statistically significant decrease in lysine in cells with the complementation plasmid (**Figure S4B**). These data suggest that the defect observed for Δ*cadA* early during acute UTI is attributed to a defect in intracellular survival.

### Lysine decarboxylation facilitates UPEC survival in the lysosomal compartment

UPEC enter the bladder epithelial cell in RAB27b^+^ fusiform vesicles (34, 43), leading to activation of Toll-like receptor 4 (TLR4), an increase in intracellular cyclic AMP (cAMP) levels and subsequent expulsion of UPEC-containing RAB27b^+^ vesicles (43). Intracellular UPEC that escape the RAB27b^+^ vacuole are targeted by autophagy and delivered into the lysosomes that have a lower pH (44). UPEC neutralize these lysosomes via a yet uncharacterized mechanism (17). To determine if the intracellular defect observed in the bladder cells is due to increased susceptibility to the low pH of the lysosome, we performed modified adherence and invasion assays using chloroquine treatment. Given that the Δ*cadA* and Δ*cadA*Δ*speF* mutants exhibited the same phenotypes (**Figure 1C, Figure 3C**), we used only Δ*cadA*Δ*speF* for this assay. Chloroquine is a lysosomotropic drug that accumulates in the acidic endosomes and not in the cytosol (45). Chloroquine has been shown to be bactericidal by inhibiting DNA and RNA synthesis (46). Previous work successfully used chloroquine to evaluate cytoplasmic versus vacuolar UPEC (47-49). This experiment revealed that while a difference in intracellular titers was observed for Δ*cadA*Δ*speF* (**Figure 4A**) as we observed previously with gentamicin treatment **(Figure 3C)**, the gentamicin/chloroquine treated samples revealed no difference in cytoplasmic CFUs **(Figure 4A)**. These data suggest that the intracellular titer defect in the mutants lacking AR4 likely stems due to lower numbers in the lysosomal vacuole.

**Figure 4.**
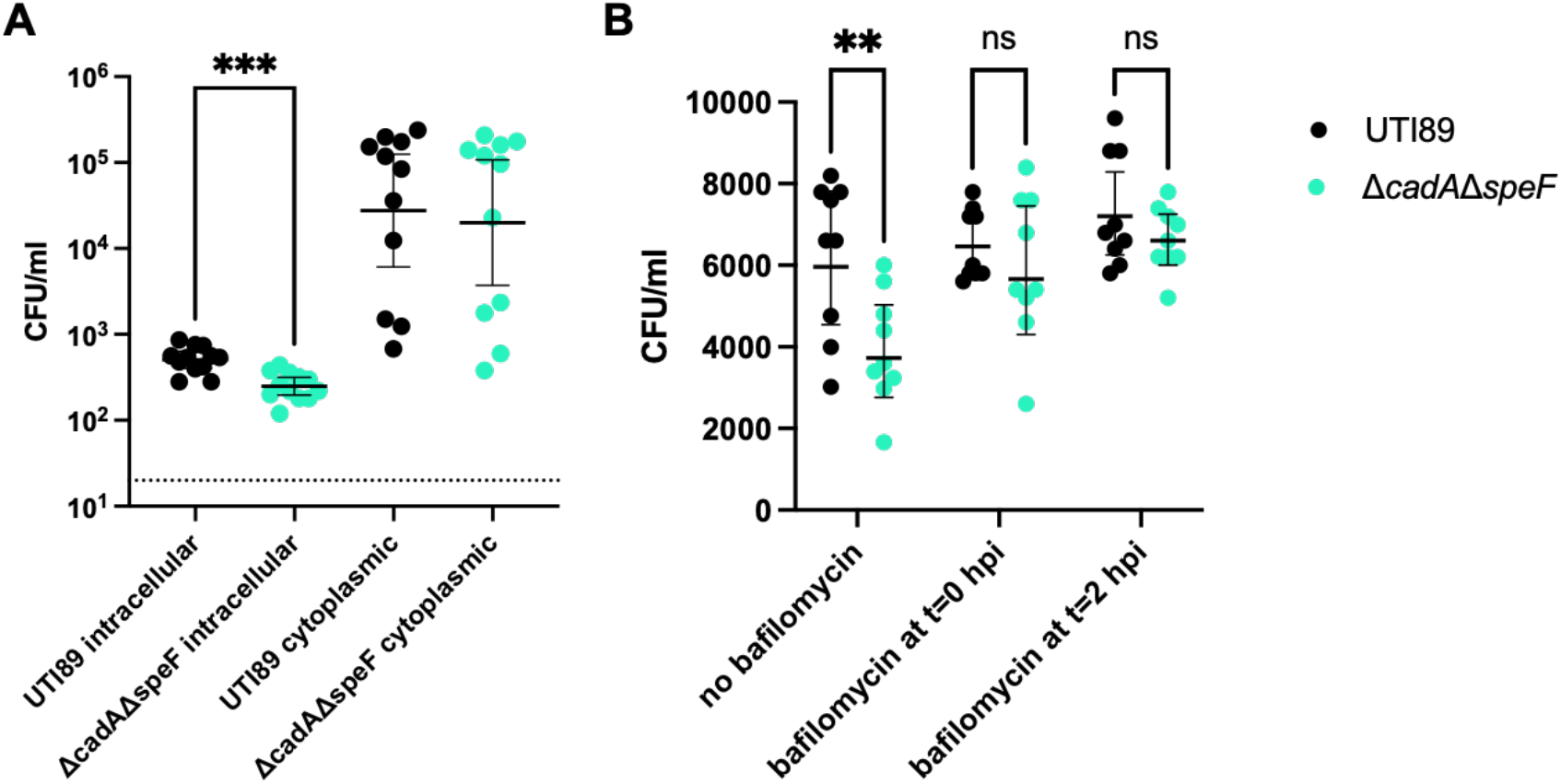
AR4 mediates survival inside the acidic vacuole. **A)** Graph depicts cytoplasmic bacterial burdens following treatment with gentamicin alone, or with gentamicin and chloroquine. Statistical test used: Kruskal Wallis with Dunn’s multiple comparisons. **B)** Graph depicts intracellular burdens in untreated urothelial cells, or in cells treated with bafilomycin at the start of infection (t=0), or at 2h following infection. **p=0.0036, two-way ANOVA with Sidak’s multiple comparisons test.

If the lack of AR4 leads to more sensitivity in the lysosomal vacuole, we reasoned that neutralization of the acidic pH may reverse the phenotype of the AR4 mutant. Bafilomycin is an inhibitor of phagosome-lysosome fusion by targeting the V-ATPase to inhibit lysosomal acidification (50). We therefore used bafilomycin to determine whether inhibition of lysosomal acidification would rescue the Δ*cadA*Δ*speF* phenotype. Indeed, addition of bafilomycin at the time of infection or at the time of gentamicin treatment restored intracellular survival of the Δ*cadA*Δ*speF* mutant to wild-type levels (**Figure 4B**). Together, these data demonstrate that AR4 aids in UPEC survival in the urothelial cell lysosome.

## Discussion

In this study, we begin to discern the distinct contributions of acid resistance/response (AR) mechanisms during UTI. Focusing on the early intracellular stage of UPEC inside urothelial cells, we present evidence that the CadA/CadBC system (AR4, **Figure 3A**) aids in UPEC survival within the bladder epithelial cell vacuoles. Thus, lysine decarboxylation contributes to the early colonization of the bladder by protecting UPEC from damaging acidic pH conditions inside the host cell. *Cad* gene expression is dependent upon both a low pH and the presence of lysine in the environment **(Figure 3A)**; here we show that AR4 is the most highly induced AR system when UPEC is inside the host cell, and we demonstrate that sufficient levels of free lysine are present in the urothelial cell to support AR4 activation. In addition to CadC sensing low pH (51), low pH in the intracellular compartment may be sensed through the PhoQ sensor histidine kinase (52). PhoQ is a histidine kinase that in response to low pH initiates signal transduction events that lead to the repression of the pleiotropic regulator HNS and subsequent de-repression of the *cad* genes (52). In recent studies we reported that deletion of the *phoPQ* two component system results in a significant defect in early bacterial colonization in the bladder, which leads to lower incidence of chronic infection in this murine model (53). We thus propose a model in which UPEC PhoQ senses the low pH of the vacuole, leads to induction of *cadA, cadB* and *cadC*, which then import the available lysine and decarboxylate it to support UPEC survival in the lysosomal compartment.

Other organisms have been shown to rely on lysine decarboxylation, including *Salmonella enterica*. However, in contrast with what we observe for UPEC, studies in *Salmonella* indicate that lysine decarboxylation genes are downregulated in the *Salmonella* containing vacuole (SCV) via the action of the OmpR response regulator (54). Notably, neutralization of the pH in the SCV inhibits secretion of the *Salmonella* pathogenicity island (SPI)-2 effector proteins that are essential for bacterial replication in the vacuole (55). As such, UPEC and Salmonella vary in that *Salmonella* which resides and replicates in the SCV suppresses AR4, whereas UPEC - which must exit the vacuole in which it is enclosed – upregulates AR4.

Defects in lysine decarboxylation have previously been shown to impair UPEC survival under nitrosative stress conditions (56, 57). However, a mechanism for cadaverine-mediated protection to nitrosative stress was not determined (56, 57). Our study builds upon the prior findings, demonstrating that AR4 deletion impacts intracellular bacterial burden, by impairing UPEC survival in the acidic vacuole of bladder epithelial cells. It is possible that neutralization of the lysosomal compartment via the decarboxylation of lysine allows for UPEC survival, affording the pathogen time to escape into the cytoplasm. It is also likely that the production of cadaverine contributes to UPEC survival. Cadaverine, like other polyamines, has been shown to influence many cellular processes. Due to its polycationic properties, it can regulate DNA, RNA, protein, and phospholipid synthesis (58). As such, polyamines have been characterized in many contrasting physiologic phenomena. In some cases, polyamines have also been shown to improve gastrointestinal barrier integrity to protect host cells from infection (58). In other cases, bacterially derived polyamines increased the abundance of anti-inflammatory macrophages in the gut and prevented the formation of the NLRP3 inflammasome, leading to improved bacterial colonization of the host (59, 60). Previous work by Jeff Purkerson has shown that the AR mechanisms that produce polyamines are important for UPEC colonization of the kidneys in a murine model of kidney acidosis (61). It is possible that cadaverine production inside the bladder cell may be altering ability of bladder cells to clear bacteria.

One of the protective responses to UTI is the shedding of umbrella cells in the bladder epithelium. This response allows for the excretion of infected cells through the urine. Polymers of L-lysine have been shown to play an important role for bladder cell exfoliation when administered as a treatment (62). We measured lysine concentrations in bladder epithelial cell culture after UPEC infection. Infection with the Δ*cadA* mutant results in higher free lysine concentrations (although not statistically significant), compared to the Δ*cadA/*p*cadA* complement strain (**Figure S6**). These data suggest that UPEC is consuming lysine within the bladder cells during infection. If the difference in lysine levels that we observe is biologically significant, we posit that UPEC consumption of lysine can potentially provide protection in two ways: 1) through the decarboxylation reaction that protects UPEC from acidic environments and 2) through the prevention of poly-L-lysine induced desquamation.

We showed that in the bladder, AR4 mutants have a significant colonization defect (**Figure 1C**). Surprisingly, these mutants do not have a defect in vaginal colonization (**Figure S3B**). This could be due to several reasons: It is possible that AR4 is dispensable for survival in the bladder lumen and the vagina, with another AR mechanism, or multiple ARs contributing to colonization of the other host environments. This would be in agreement with recent data we reported, indicating niche-specific use of two-component systems in the same murine model (53). We speculate that the other AR mechanisms are not expendable but are needed at different times during infection. For example, the herein observation of Δ*gadA*Δ*gadB* exhibiting a defect in the bladder may indicate a role of AR2 in UPEC survival in the lumen. Likewise, our data point towards a potential role of AR3 in colonization of the kidney (**Figure S3A**) Collectively, our work begins to elucidate how UPEC responds to acidic conditions during UTI. We provide, for the first time, evidence of a specific AR mechanism being critical for the survival of UPEC in the urothelial cell acidic vacuole.

## Materials and Methods

### Bacterial Strains and Growth Conditions

All studies were performed in the well characterized UPEC cystitis isolate UTI89 and derived isogenic deletion mutants. UTI89 is of the sequence type ST95 and is serotyped as O18:K1:H7 (28). Strains were propagated in unbuffered lysogeny broth (LB) (Fisher Scientific), at pH 7.4. Inoculated strains were grown overnight at 37ºC with shaking unless otherwise noted. Gene deletions were created using the λ-red recombinase system (63). A complete list of strains, plasmids, and primers used in this study can be found in **Table S1 and S2** respectively.

For growth analyses in pooled urine, urine was collected from healthy human volunteers in accordance with approved protocols (IRB #151465). A healthy volunteer is defined as an individual who is urologically asymptomatic, not menstruating, and who has not taken antibiotics in the last 90 days. An equal volume of male and female urine was pooled from multiple volunteers then filtered through a 0.22 μm filter before use.

### Cell Lines

Cell culture infections were performed using 5637 human transitional bladder epithelial cells (ATCC HTB-9) originally derived from a 68-year old white male with Grade II bladder carcinoma. 5637 cells were propagated and infected in RPMI 1640 (Gibco) supplemented with 10% fetal bovine serum (Gibco).

### Adherence and invasion assay

HTB-9 bladder epithelial cells were seeded at 1.0×10^5^ cells/well in 24 well plates. Once cells reach at least 80% confluence, they were infected with UTI89 or the UTI89 acid resistance mutants at an MOI of 7.5. The plates were then centrifuged at 600xg for 5 minutes to facilitate and synchronize attachment. The cells were then incubated for 2 hours at 37ºC in 5% CO_2_. To quantify total CFUs, monolayers were treated with 0.1% Triton X-100 and serially diluted to enumerate CFUs/ml. To determine adherent CFUs, monolayers were washed with PBS three times to remove extracellular bacteria, lysed with 0.1% Triton X-100 and serially diluted. To determine intracellular CFUs, monolayers were washed with PBS three times then treated with PBS containing 100 ug/ml gentamicin to kill extracellular bacteria. After incubation in gentamicin for 2 hours, the monolayers were washed three times in PBS then lysed with 0.1% Triton X-100 and serially diluted to enumerate CFUs/ml.

### Murine infections

Murine infections were performed as described previously (64). Briefly, UTI89 and each mutant strain were inoculated into 5 mL of LB and grown shaking at 37C overnight. Next, the overnight cultures were diluted 1:1000 in 10 mL of fresh LB and grown statically at 37ºC for 24 hours. These static cultures were then diluted 1:1000 in 10 mL of fresh LB and grown statically at 37C for another 24 hours. Next, each culture was normalized to 10^7^ CFU in PBS. 7-week-old C3H/HeN female mice (Envigo) were transurethrally inoculated with 50 μl of the normalized bacteria in PBS. Mice were sacrificed at 24 hours post infection. The bladders, kidneys, and vaginas were harvested and homogenized for CFU enumeration. All animal studies were approved by the Vanderbilt University Medical Center Institutional Animal Care and Use Committee (IACUC) (protocol number #M1800101-01) and carried out in accordance with all recommendations in the Guide for the Care and Use of Laboratory Animals of the National Institutes of Health and the IACUC.

### Intracellular Free Lysine Quantitation

Using a targeted metabolomics method to measure intracellular free lysine, 3×10^6^ cells were seeded in T75 flasks. The next day, media was removed, flasks were washed twice with room temperature PBS and cells were scraped into PBS to harvest. The cells were then pelleted in 15 mL conical tubes, flash frozen in liquid N_2_, and stored at -80°C until ready for metabolite extraction. 30 nmol of internal standard (^13^C_6_,^15^N_2_-Lysine, Thermo Fisher Scientific, Waltham, MA) was spiked into each sample. Metabolites were extracted by adding 2 mL of -80°C 80:20 MeOH:H_2_O to the cell pellets, vortexed vigorously, and then placed in a -80°C freezer to extract for 15 minutes. After 15 minutes, samples were pelleted by centrifugation at 3,500 x *rpm* for 10 minutes at 4°C. Supernatant containing extracted metabolites and internal standard was transferred to new 15 mL conical tubes and dried under N_2_. Precipitated protein pellets were resolubilized and protein concentration was measured via BCA assay (Thermo Fisher Scientific, Waltham, MA). Samples were resuspended in 80 μL 3:2 mobile buffer A: mobile buffer B (see below). 18 μL of the sample was then chromatographed with a Shimadzu LC system equipped with a 100 × 2.1mm, 3.5μm particle diameter XBridge Amide column (Waters, Milford, MA). Mobile phase A: 20 mM NH_4_OAc, 20 mM NH_4_OH, 5% acetonitrile in H_2_O, pH 9.45; mobile phase B: 100% acetonitrile. With a flow rate of 0.45 mL/min the following gradient was used: 2.0 min, 95% B; 3.0 min, 85% B; 5.0 min, 85% B; 6.0 min, 80% B; 8.0 min, 80% B; 9.0 min, 75% B; 10 min, 75% B; 11 min, 70% B; 12 min, 70% B; 13 min, 50% B; 15 min, 50% B; 16 min 0% B; 17.5 min, 0% B; 18 min, 95% B. The column was equilibrated for 3 minutes at 95% B between each sample. Scheduled MRM was conducted in negative mode with a detection window of 120 seconds using an AB SCIEX 6500 QTRAP with the analyte parameters below. Lysine was then quantified via LC-MS/MS using the ^13^C_6_,^15^N_2_-Lysine internal standard and normalized to the protein in each respective sample’s cell pellet. Outliers were removed using interquartile range.

| Species | Q1 (m/z) | Q3 (m/z) | RT (min) |
| --- | --- | --- | --- |
| Lysine | 147 | 84 | 14 |
| $^{13}\text{C}_6, ^{15}\text{N}_2$-Lysine [Internal Standard] | 155 | 90 | 14 |

## Acknowledgments

This research was supported by grants: R01AI168468 (to MH), 2T32AI112541-06 (to MAW), T32 GM007569 (to JRB), F30AI169748 (to SAR), F31AI174488 (to TAB).

## Author contributions

MAW and MH conceived the study, performed the experiments, analyzed the data, and composed the manuscript. SR, TB, GM, JRB, FG, and EQJ performed experiments and edited the manuscript. JR provided access to the instrumentation needed for lysine quantification.

