## SupplementaryFiguresAndTables for "Lysine Decarboxylation aids in UPEC intracellular survival in the early stages of urinary tract infection"

### Supplementary Figures

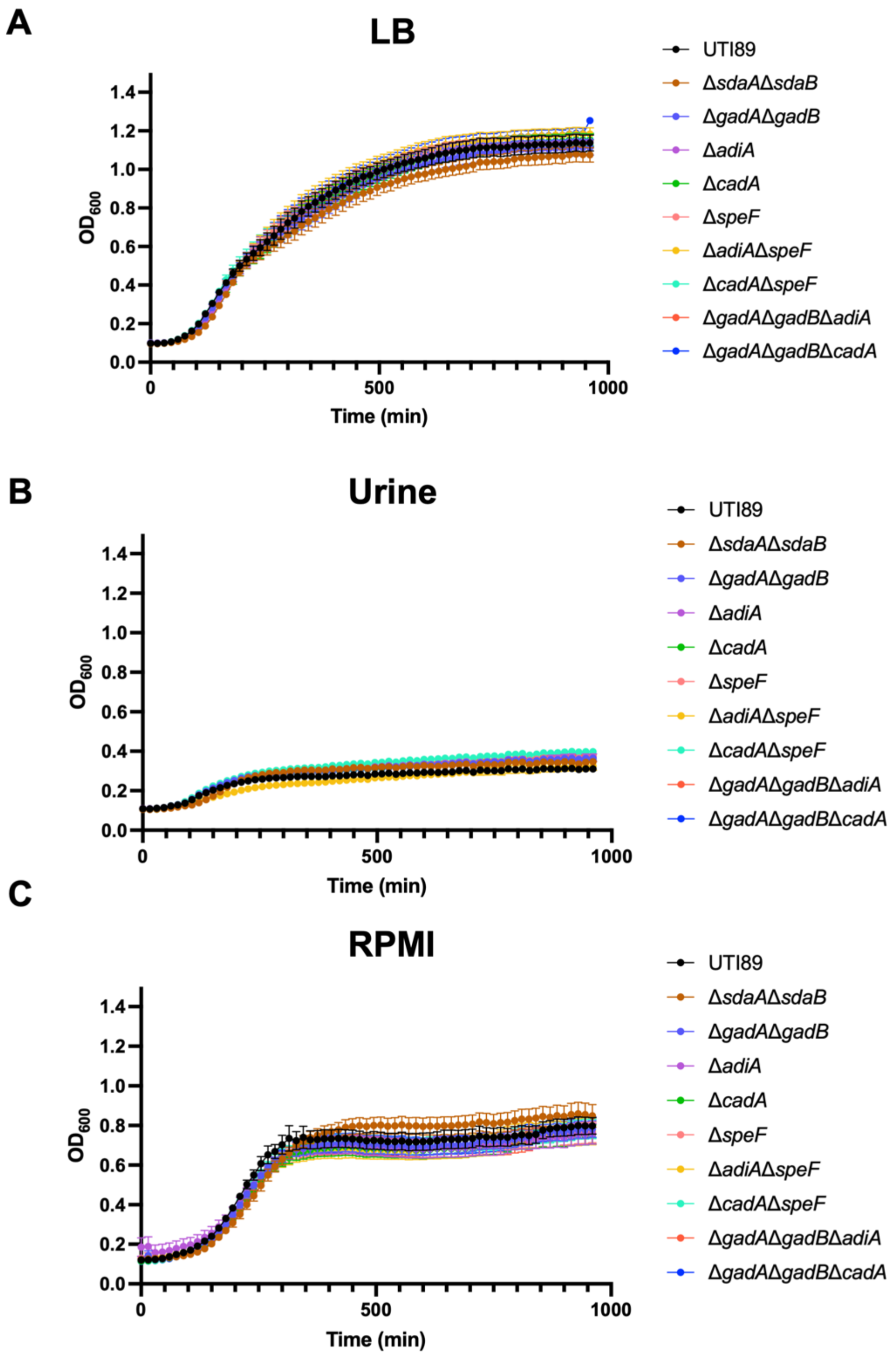

**Figure S1. Growth curves of strain UTI89 and isogenic acid resistance mutants.** UTI89 and the indicated AR mutants were grown in LB (A), pooled human urine (B), and RPMI (C) in 96 well plates for 16 hours. $OD_{600}$ was measured using the plate reader every 15 minutes. Graphs depict 2 biological replicates.

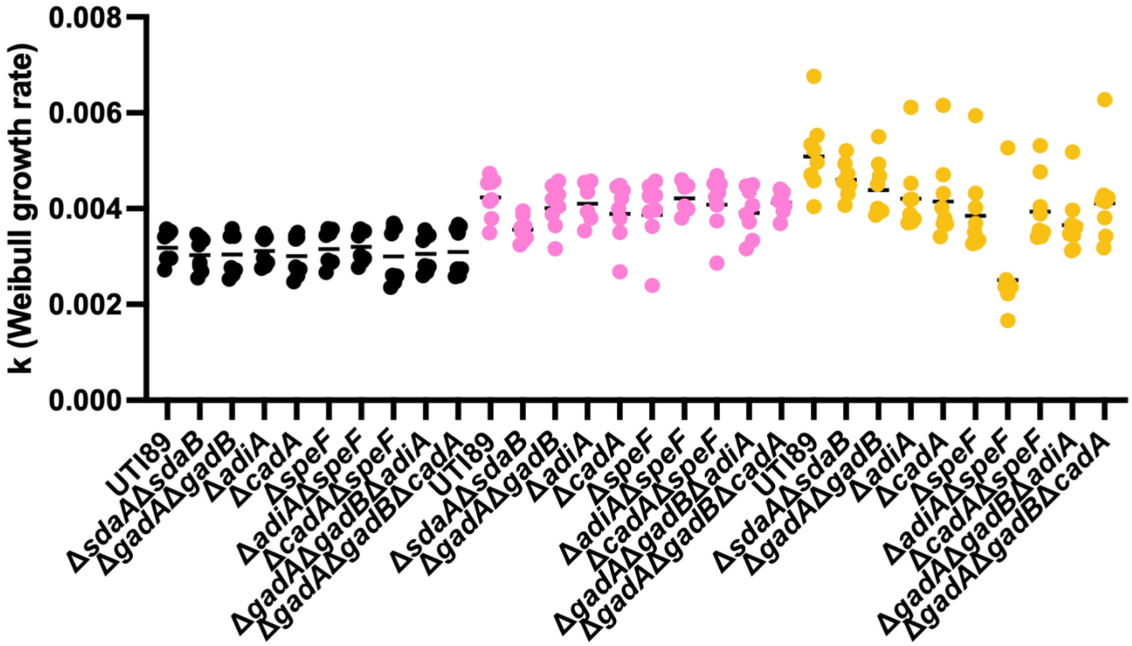


**Figure S2. Growth rates of AR mutants.** Growth rates for cells grown in LB (black), RPMI (pink), or pooled human urine (yellow) were calculated using the Weibull growth curve formula.

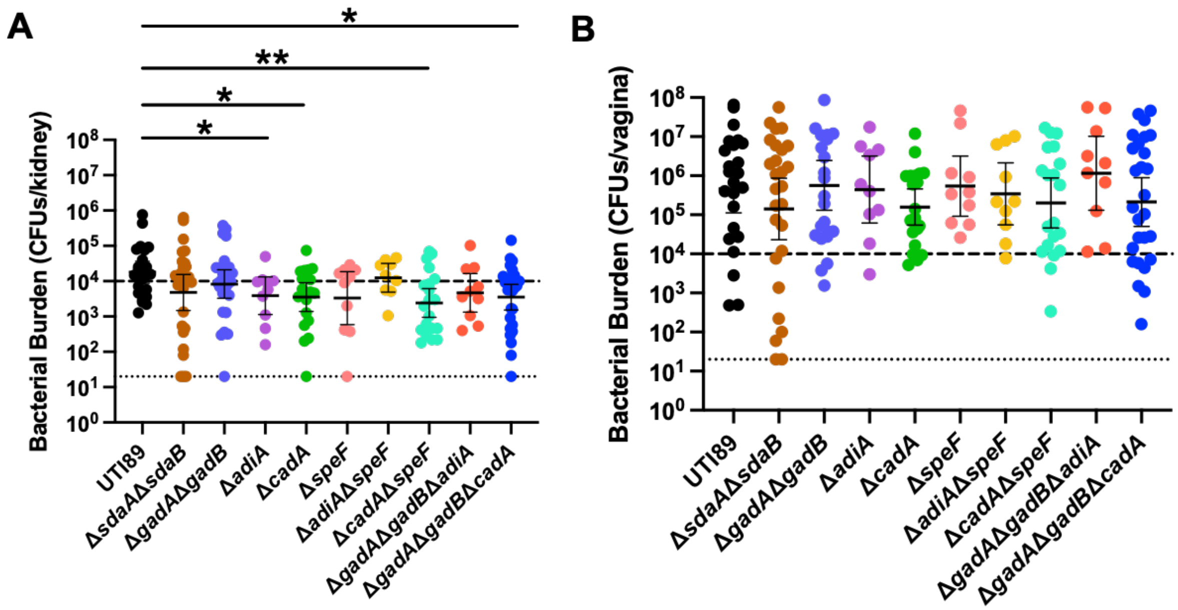


**Figure S3. Fitness of AR deletion mutants in the kidney and vaginal space during acute UTI.** Graphs depict bacterial titers in the kidneys (A) and vaginal membranes (B) of mice 24h after transurethral inoculation with wild-type UTI89 or the isogenic AR mutants. Kidneys and vaginas were harvested, homogenized, and plated for CFU enumeration. *p<0.05, **p<0.002, by two-tailed Mann-Whitney.

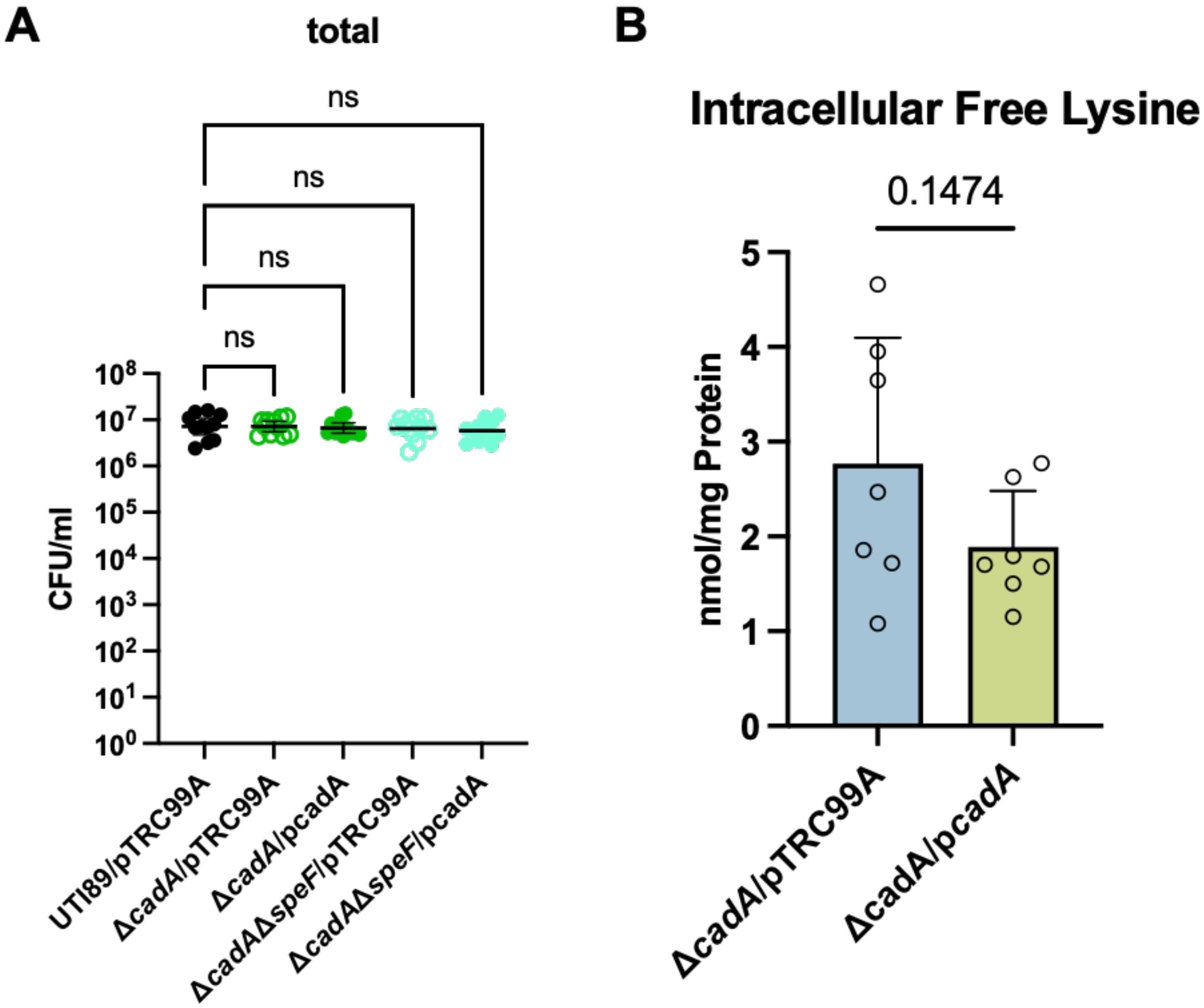


**Figure S4. Effect of *cadA* deletion on bacterial survival and lysine abundance inside the urothelial cells. A)** Graph depicts total CFUs harnessed from HTB-9 cells infected with UTI89 (black), Δ*cadA* (green), or Δ*cadA*Δ*speF* (cyan) at an MOI of 7.5. Total CFUs were enumerated at 4h post inoculation. Statistical test performed: One-Way ANOVA with Dunnett's multiple comparisons. **B)** Bar graph shows the concentration of free intracellular lysine in HTB9 bladder cells infected with Δ*cadA*/pTRC99A or Δ*cadA*/pcadA, at 4h post inoculation, following a 2-hour gentamicin treatment (n=7 biological replicates, One-Way ANOVA with Tukey multiple comparisons test).

**Table S1. Strains and plasmids used in this study**

| *E. coli* strain | Relevant genotype | Plasmid |
|---|---|---|
| UTI89 | | N/A |
| UTI89 Δ*sdaA*Δ*sdaB* | Δ*sdaA*Δ*sdaB*[1] | N/A |
| UTI89 Δ*gadA*Δ*gadB* | Δ*gadA*Δ*gadB*[1] | N/A |
| UTI89 Δ*adiA* | Δ*adiA*[1] | N/A |
| UTI89 Δ*cadA* | Δ*cadA*[1] | N/A |
| UTI89 Δ*speF* | Δ*speF*[1] | N/A |
| UTI89 Δ*cadA*Δ*speF* | Δ*cadA*Δ*speF* | N/A |
| UTI89 Δ*gadA*Δ*gadB*Δ*cadA* | Δ*gadA*Δ*gadB*Δ*cadA* | N/A |
| UTI89 Δ*gadA*Δ*gadB*Δ*adiA* | Δ*gadA*Δ*gadB*Δ*adiA* | N/A |
| UTI89/pTRC99A | Empty vector pTRC99A | pTRC99A |
| UTI89 Δ*cadA*/pTRC99A | Δ*cadA* with empty vector pTRC99A | pTRC99A |
| UTI89 Δ*cadA*/p*cadA* | Δ*cadA* with *cadA* complemented on the pTRC99A plasmid | pTRC99A |
| UTI89 Δ*cadA*Δ*speF*/pTRC99A | Δ*cadA*Δ*speF* with empty vector pTRC99A | pTRC99A |
| UTI89 Δ*cadA*Δ*speF*/p*cadA* | Δ*cadA*Δ*speF* with *cadA* complemented on the pTRC99A plasmid | pTRC99A |

1. Wiebe Michelle, A., Brannon John, R., Steiner Bradley, D., Bamidele, A., Schrimpe-Rutledge Alexandra, C., Codreanu Simona, G., Sherrod Stacy, D., McLean John, A., and Hadjifrangiskou, M. (2022). Serine Deamination Is a New Acid Tolerance Mechanism Observed in Uropathogenic Escherichia coli. mBio *13*, e02963-02922. 10.1128/mbio.02963-22.

**Table S2. Primers and probes used in this study**

| Primer name | Sequence 5'-3' | Purpose |
|---|---|---|
| sdaA_KO_f | GTTATTAGTTCGTTACTGGAAGTCCAGTCACCTTGTCAGGAGTATTATCGTGTAGGCTGGAGCTGCTTC | Deletion of *sdaA* |
| sdaA_KO_r | AAGCGGGAATAAATTCGCCCATCCGTTGCAGATGGGCGAATAAGAAGATCATATGAATATCCTCCTTAG | Deletion of *sdaA* |
| sdaA_KO_test_f | CATCTGGGTCGTTATCATCCT | Validation of *sdaA* deletion |
| sdaA_KO_test_r | GTAACGAGTGCGCAAATCG | Validation of *sdaA* deletion |
| sdaB_KO_f | CGCGCCGCTTTCGGGCGGCGCTTCCTCCGTTTTAACGCGATGTATTTCCTGTGTAGGCTGGAGCTGCTTC | Deletion of *sdaB* |
| sdaB_KO_r | GGATGAGAAATCGGGAAGAGGCCTCGCAAAAAGAGGCCTCTGGAGAGCGACATATGAATATCCTCCTTAG | Deletion of *sdaB* |
| sdaB_KO_test_f | GTTCCTGATGCCGATGTAC | Validation of *sdaB* deletion |
| sdaB_KO_test_r | CCAGAACAGGCTATGGCT | Validation of *sdaB* deletion |
| sdaC_KO_f | GGCTGAACTGGCTAAAAGCTGAATTATTTGCATTCCTCCAGGAGAAATAGGTGTAGGCTGGAGCTGCTTC | Deletion of *sdaC* |
| sdaC_KO_r | ACATCGCGTTAAAACGGAGGAAGCGCCGCCCGAAAGCGGCGCGAAAGGACCATATGAATATCCTCCTTAG | Deletion of *sdaC* |
| sdaC_KO_test_f | CATCGCCGATAGACAGAT | Validation of *sdaC* deletion |
| sdaC_KO_test_r | GAACTCCACTTCATGCTGAC | Validation of *sdaC* deletion |
| gadB_KO_FOR | CAGGTGTGTTTAAAGCTGTTCTGCTGGGCAATACCCTGCAGTTTCGGGTGTGTAGGCTGGAGCTGCTTC | Deletion of *gadB* |
| gadB_KO_REV | CAAGTAACGGATTTAAGGTCGGAACTACTCGATTCACGTTTTGGTGCGAACATATGAATATCCTCCTTAG | Deletion of *gadB* |
| gadB_KO_Test_FOR | GTGAACAGACTTTGGAAATTGTCCC | Validation of *gadB* deletion |

| | | |
|---|---|---|
| gadB_KO_Test_REV | ACTTGCTTACTTTATCGATAAATCCTA | Validation of *gadB* deletion |
| gadA_KO_FOR | GTTTAAAGCTGTTCTGCTGGGCAATACCCTGCAGTTTCGGGTGGTCGCTGGTGTAGGCTGGAGCTGCTTC | Deletion of *gadA* |
| gadA_KO_REV | AAATGGACCAGAAGCTGTTAACGGATTTCCGCTCAGAACTACTCGATTCACATATGAATATCCTCCTTAG | Deletion of *gadA* |
| gadA_KO_Test_FOR | CAATTAATAAGTAGCCGAATACCCACC | Validation of *gadA* deletion |
| gadA_KO_Test_REV | TGTAATACCTTGCTTCCATTGCG | Validation of *gadA* deletion |
| adiA_ko_f | ATGATGAAAGTATTAATTGTTGAAAGCGAGTTTCTCCATCAAGACACCTGGTGTAGGCTGGAGCTGCTTC | Deletion of *adiA* |
| adiA_ko_r | TTACGCTTTCACACACATAACGTGGTAAATACCGTCAATAATTTCTGTCCCTTCCATATGAATATCCTCCTTAG | Deletion of *adiA* |
| adiA_kotest_f | GAAGATACTTGCCCGCAAC | Validation of *adiA* deletion |
| adiA_kotest_r | CTCGCTAAAGCGAAGCGATAC | Validation of *adiA* deletion |
| cadA_ko_f | ATGACTATGAACGTTATTGCAATATTGAATCACATGGGGGTTTATTTTAAAGAAGGTGTAGGCTGGAGCTGCTTC | Deletion of *cadA* |
| cadA_ko_r | TTATTTTTTGCTTTCTTCTTTCAATACCTTAACGGTATAGCGGCCATCAGCATATGAATATCCTCCTTAG | Deletion of *cadA* |
| cadA_kotest_f | GTACCTTCATCGTCAGCCTG | Validation of *cadA* deletion |
| cadA_kotest_r | GTGTTCTCCTTATGAGC | Validation of *cadA* deletion |
| speF_ko_f | ATGACGAGTATAGCCAGTTACCGGGCTGGTCTGGGTTATTGCATCTGCGTGTAGGCTGGAGCTGCTTC | Deletion of *speF* |
| speF_ko_r | AATTTTTCCCCTTTCAACAGGGCGCTTTGCGCATCACGAGGCTTGATGACCATATGAATATCCTCCTTAG | Deletion of *speF* |
| speF_kotest_f | GGTGCTCATATACTGCTAAC | Validation of *speF* deletion |
| speF_kotest_r | GTTGACCATCGTCAGTATG | Validation of *speF* deletion |

| | | |
|---|---|---|
| cadA_pTRCinsert_F | atttcacacaggaaacagaccatggTTATTTTTTGCTTTCTTCTTTCAATACC | Amplification of *cadA* for insertion into pTRC |
| cadA_pTRCinsert_R | ctcatccgccaaaacagccaagcttCGGTAACTTCCCGAAAGTTTATG | Amplification of *cadA* for insertion into pTRC99A |
| pTRC_linearization_F | AAGCTTGGCTGTTTTGGC | Linearization of pTRC99A |
| pTRC_linearization_R | CCATGGTCTGTTTCCTGTG | Linearization of pTRC99A |
| adiA_qPCR_F | CTGGTCGGTAGTCGTCGGTA | *adiA* qPCR |
| adiA_qPCR_R | GCCTGTCAGCATCAAGCCTTG | *adiA* qPCR |
| adiA_qPCR_probe | FAM-CGATAACGATGTCGTGGTCGTTGACCGTAACTG-MGBNFQ | *adiA* qPCR |
| cadA_qPCR_F | GTGCTGCTGATGATATTGCTAAC | *cadA* qPCR |
| cadA_qPCR_R | GAATGCAGTACCACCCATG | *cadA* qPCR |
| cadA_qPCR_probe | FAM-CGAATATATCAACACTATTCTGCCTCCGCTGA-MGBNFQ | *cadA* qPCR |
| gadA_qPCR_F | GATGTGGAGTTGCGTGAGATC | *gadA* qPCR |
| gadA_qPCR_R | CTCATAGTTACCGGTGTAGGTC | *gadA* qPCR |
| gadA_qPCR_probe | FAM-ACCCGAAACGCATGATTGAAGCCTGCGACG-MGBNFQ | *gadA* qPCR |
| gadC_qPCR_F | CGTTGAAGCTTCCGCAACCCA | *gadC* qPCR |
| gadC_qPCR_R | CTGCGGAGAGGTTGATTTCATTAC | *gadC* qPCR |
| gadC_qPCR_probe | FAM-TCCACTGGCTATGTTACTGCTGATGGTGGCGG-MGBNFQ | *gadC* qPCR |
| sdaA_qPCR_f | TGCAAATCCACGCCTATAACG | *sdaA* qPCR |
| sdaA_qPCR_r | CAGTACGCGAGCAGTTCG | *sdaA* qPCR |
| sdaA_qPCR_probe | FAM-CGAAGTGAGCGTGCCGTATCCG-MGBNFQ | *sdaA* qPCR |
| sdaC_qPCR_f | GATGCTGCTGGCTCTGTACC | *sdaC* qPCR |

| | | |
|---|---|---|
| sdaC_qPCR_r | ATGATCGGAGAGTGGTTGAACG | *sdaC* qPCR |
| sdaC_qPCR_probe | NED-GCTGTCTCTGGACACTGCATCTG-MGBNFQ | *sdaC* qPCR |
| speF_qPCR_F | GAAGGTGTCAGCGGTCGTA | *speF* qPCR |
| speF_qPCR_R | GCTGTTTCATACGACTGCCAG | *speF* qPCR |
| speF_qPCR_probe | FAM-ATGAACAGTTTATTCCGATGATGGCGGACTG-MGBNFQ | *speF* qPCR |
| gyrB_qPCR_f | GATGCGCGTGAAGGCCTGATTG | qPCR housekeeping gene |
| gyrB_qPCR_r | CACGGGCACGGGCAGCATC | qPCR housekeeping gene |
| gyrB_qPCR_probe | VIC-ACGAACTGCTGGCGGA-MGBNFQ | qPCR housekeeping gene |
